# The dynamical Matryoshka model: 3. Diffusive nature of the atomic motions contained in a new dynamical model for deciphering local lipid dynamics

**DOI:** 10.1101/2022.03.30.486412

**Authors:** Tatsuhito Matsuo, Aline Cisse, Marie Plazanet, Francesca Natali, Michael Marek Koza, Jacques Ollivier, Dominique J. Bicout, Judith Peters

## Abstract

In accompanying papers [Bicout et al., BBA – Biomembr. *Submitted*; Cissé et al., BBA – Biomembr. *Submitted*], a new model called Matryoshka model has been proposed to describe the geometry of atomic motions in phospholipid molecules in bilayers and multilamellar vesicles based on their quasielastic neutron scattering (QENS) spectra. Here, in order to characterize the relaxational aspects of this model, the energy widths of the QENS spectra of the samples were analyzed first in a model-free way. The spectra were decomposed into three Lorentzian functions, which are classified as slow, intermediate, and fast motions depending on their widths. The analysis provides the diffusion coefficients, residence times, and geometrical parameters for the three classes of motions. The results corroborate the parameter values such as the amplitudes and the mobile fractions of atomic motions obtained by the application of the Matryoshka model to the same samples. Since the current analysis was carried out independently of the development of the Matryoshka model, the present results enhance the validity of the model. The model will serve as a powerful tool to decipher the dynamics of lipid molecules not only in model systems, but also in more complex systems such as mixtures of different kinds of lipids or natural cell membranes.

## 1. Introduction

Cell membranes play an essential role in cell function, including the boundary formation of organelles and cells, and the transport of chemical substances exploiting their permeability. It is known that cell membranes consist of various types of lipids and alter their lipid composition to adapt to extracellular environments such as temperature and pH so that they function properly at given environments [1,2]. It is thus important to understand the physical and chemical properties of lipid molecules in detail to reveal the mechanism of their biological functions. Lipid membranes can undergo phase transitions from crystalline, gel, to fluid phases depending on temperature or pressure, which impact their dynamical properties as well as their structures. The motions within membranes span a wide range of time- and space-scales: The intramolecular motions, including the rotation of methylene groups, occur in the picosecond regime, while large-scale motions, such as membrane undulations, take place from the nanosecond to the millisecond regimes [3–7]. In order to understand their dynamical nature in different physicochemical environments, various experimental techniques have been used such as incoherent neutron scattering [8], nuclear magnetic resonance spectroscopy (NMR) [9], fluorescence recovery after photobleaching (FRAP) [10] depending on the time- and space-scale of interest. Together with the advancement in molecular dynamics (MD) simulation techniques and their combination with experiments [11–13], a huge amount of knowledge of individual and collective lipid motions has been accumulated. However, there does not still exist a unified dynamical framework that describes comprehensively the lipid motions occurring at different time domains.

Among the experimental techniques employed so far, quasielastic neutron scattering (QENS) is a powerful tool to study the dynamics of lipid molecules at the pico-nanosecond timescale and Å–nm length scale, which correspond to the crossover region from localized vibrations to various diffusional processes occurring at a much longer timescale involving bilayer bending and membrane undulation. In particular, QENS provides detailed information on the motions of hydrogen atoms since the incoherent scattering cross-section of the hydrogen atom is much larger than that of any other atom found in biomolecules or its isotope deuterium [14]. Moreover, another advantage of this technique is that various motions which occur at different time scales can be focused on by changing the energy resolution of the neutron spectrometers, which is equal to shifting the time window of the observation.

In order to interpret the QENS spectra, thereby capturing the dynamical properties of lipid molecules in detail, establishment of a theoretical framework that describes relevant atomic motions is indispensable. In 1989, the first theoretical model describing the lipid motions in bilayers was reported [4], where different motional processes were assumed depending on the energy resolutions and the sample conditions. Similar approaches have widely been employed not only for lipid molecules [5,15–17], but also for other molecules such as cholesterol inserted to lipid bilayers [18]. Later, a more sophisticated model has been presented, where QENS spectra were decomposed into three kinds of motions with distinct time scales, i.e. slow, intermediate, and fast motions [19,20]. In this model, each component of these three types of motions was interpreted by various combinations of motions in lipid molecules. Although all of these studies have significantly advanced our understanding of the dynamics of lipid molecules and helped provide a physical picture, a more holistic description of the possible motions contained in the lipid molecules was still missing.

Based on the dynamical models above, a new theoretical model named Matryoshka model has been presented in the accompanying papers [21,22], where the dynamic structure factor S(Q, ω) is analytically expressed by directly incorporating the scattering functions of various motions that lipid molecules undergo in addition to the decomposition of all the possible motions into three classes, i.e. slow, intermediate, and fast motions. This model successfully accounts for the elastic incoherent structure factor (EISF) and the quasi-elastic incoherent structure factor (QISF) profiles, which are derived from the geometry of atomic motions in the samples, of 1,2-dimyristoyl-*sn*-glycero-3-phosphocholine (DMPC) in multi-lamellar bilayers (MLB) and multi-lamellar vesicles (MLV), and 1,2-dimyristoyl-*sn*-glycero-3-phospho-(1’-rac-glycerol) (DMPG) in MLB at various temperatures. This has enabled the detailed investigation of the geometry of motions in lipid molecules placed in various physicochemical environments. On the other hand, although the diffusive nature of the motions classified as the slow, intermediate and fast motions in the Matryoshka model is theoretically characterized through a series of analytical equations of relaxation rates of each of these motions [21], it has not yet been experimentally characterized. In the present paper, this aspect of each of the three types of motions are reported and discussed.

## 2. Materials and methods

### 2-1. Sample preparation and neutron scattering experiments

Details of sample preparations, QENS experiments, and the analysis of the QENS spectra are described in the accompanying papers [21,22]. Briefly, the following samples were used for neutron experiments:

- DMPC in MLB oriented at 135° relative to the incident neutron beam (DMPC MLB135), which mainly focuses on the in-plane motions
- DMPC in MLB oriented at 45° to focus on the out-of-plane motions (DMPC MLB45)
- DMPC in MLB oriented at 135° with hydrogen atoms in the tails perdeuterated (d54DMPC)
- DMPC in MLV (DMPC MLV)
- DMPG in MLB oriented at 135° relative to the incident neutron beam (DMPG MLB135), focusing on the in-plane motions

All the samples except for DMPG MLB135 were measured on the IN6 spectrometer (https://www.ill.eu/users/instruments/instruments-list/sharp/description/instrument-layout) at the Institut Laue-Langevin (ILL; in Grenoble, France) with the energy resolution of ~70 μeV, which corresponds to a time window of ~20 ps. The spectra of DMPC MLB135 and DMPG MLB135 were measured on the IN5 spectrometer (https://www.ill.eu/users/instruments/instruments-list/in5/description/instrument-layout) at the ILL with the energy resolution of ~90 μeV (~15 ps). Each spectrum was fitted by the following phenomenological equation [14] that contains three types of motions with different relaxational times:

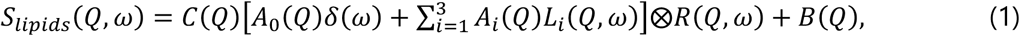

where 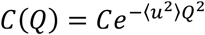 represents the scaling factor C including the Debye-Waller factor with ⟨*u*^2^⟩ the mean square amplitude of vibration, A_0_(Q)δ(ω) is an elastic component with A_0_(Q) being the elastic incoherent structure factor (EISF) and δ(ω) being the Dirac delta function. L_i_(Q, ω) (= (1/π) × (Γ_i_(Q) /(Γ_i_(Q)^2^ + ω^2^)), and i = 1, 2, 3) is the i-th Lorentzian function describing the atomic motion in lipids. Γ_i_(Q) is the half-width at half-maximum (HWHM) of L_i_(Q, ω). R(Q, ω) refers to the instrumental resolution function, which is obtained from the Vanadium measurement, B(Q) is a flat background, that includes the instrument contribution, and/or fast vibrational motions outside the current time window. Finally, ⊗ denotes the convolution operation. In the following, the three Lorentzian functions are classified as the slow motions (the narrowest Lorentzian), the intermediate motions, and the fast motions (the broadest Lorentzian). The spectra in the range of 0.37 Å^−1^ ≤ Q ≤ 2.02 Å^−1^ (for the IN6 data) and 0.33 Å^−1^ ≤ Q ≤ 1.82 Å^−1^ (for the IN5 data) were used for the fitting. Note, however, that for the spectra obtained on IN6, the HWHMs of the Lorentzian functions at the first two lowest Q values were found to show unrealistically higher values than other data points probably due to multiple scattering effects. Therefore, the spectra in the range of 0.72 Å^−1^ ≤ Q ≤ 2.02 Å^−1^ were analyzed for IN6 data and the data points at Q = 0.37 [Å^−1^] and 0.55 [Å^−1^] were excluded from the analysis and not presented in this study. Fits were carried out in the range of −10 meV ≤ ΔE ≤ 2 meV (for the IN6 data) and −10 meV ≤ ΔE ≤ 1.3 meV (for the IN5 data) using IGOR Pro software (WaveMetrics, Lake Oswego, OR, USA). Some examples of fits are shown in Fig. 1.

**Figure 1.**
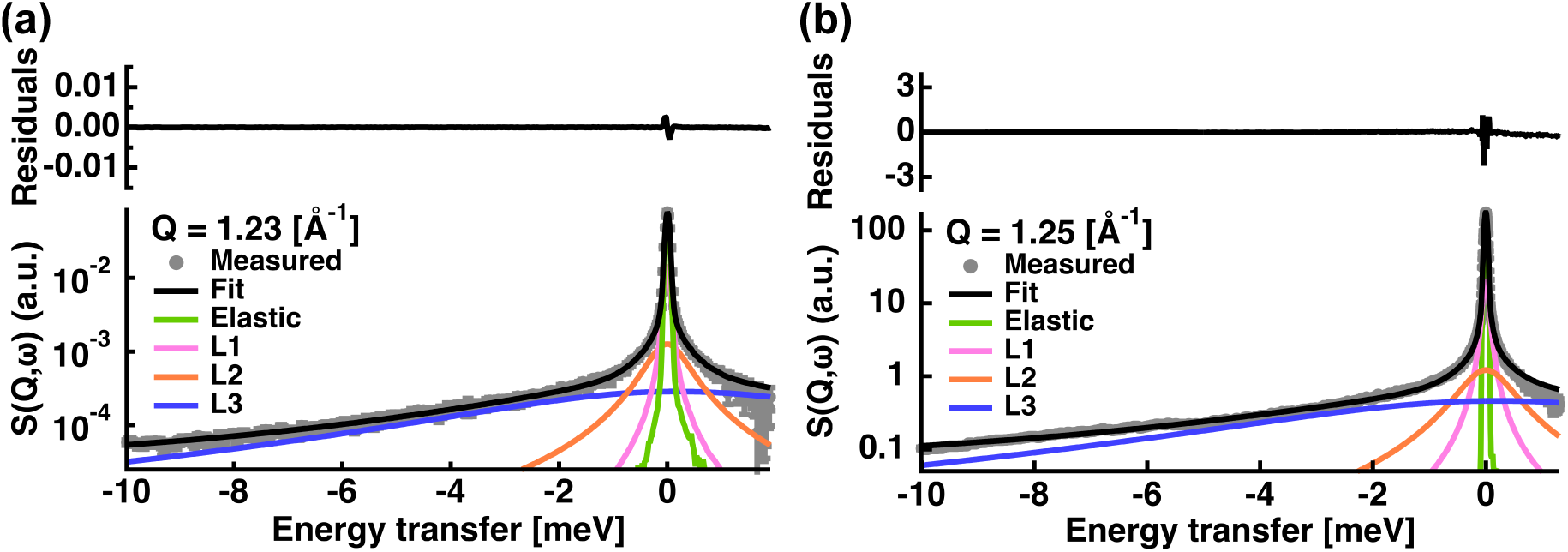
Examples of fits of the QENS spectra of DMPC MLB135 measured on IN6 (a) and IN5 (b). (a) and (b) are the data at Q = 1.23 Å^−1^ and T = 283 K, and at Q = 1.25 Å^−1^ and T = 280 K, respectively. The grey circles are the experimental data points with corresponding statistical errors. The total fit is represented by the black line. The green line corresponds to the elastic peak convoluted by the resolution function, which is directly given by the Vanadium measurements. The magenta, orange and blue curves are the Lorentzian functions convoluted by R(Q, ω) for slow, intermediate and fast motions, respectively. The upper panels show the residuals of the fits.

### 2-2. Lipid motions contained in the Matryoshka model

In the Matryoshka model [21] as shown schematically in Fig. 2, a lipid molecule is represented by a head group and a tail group (with only one tail instead of two for the sake of simplicity), which contain a fraction z and a fraction (1-z) of the total number of hydrogen atoms, respectively. The values of z are 0.25 and 0.18 for DMPC and DMPG, respectively. The slow, intermediate, and fast motions contain the following internal and/or collective motions of the lipid molecules (Table 1):

**Table 1.** Classification of various motions contained in the Matryoshka model.

|  | Internal motion | Collective motion |
| --- | --- | --- |
| Slow motions |  | 2D-diffusion<br>in-out of the plane diffusion |
| Intermediate motions | Head-flip-flop<br>Rotation (head) | Rotational diffusion |
| Fast motions | 2D-diffusion (tail)<br>Jump diffusion (tail and head) |  |

**Figure 2.**
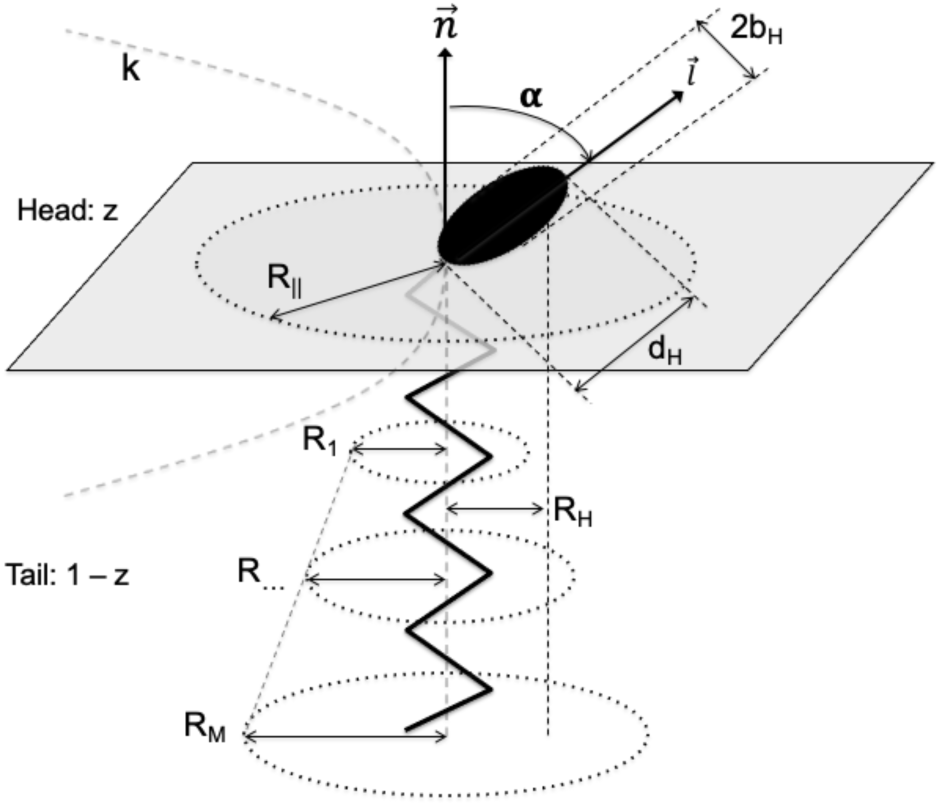
Schematic illustration of the Matryoshka model. A lipid molecule in membranes is drawn, on which parameters employed in the Matryoshka model are shown (for details of the parameters, refer to the main text). This figure is taken from Figure 2 of [21].

i. The slow motions contain the following two movements:

- A whole lipid molecule undergoes two-dimensional diffusion (2D-diffusion) within the membrane plane. The space where the lipid molecule can move is represented by a cylinder with the radius of R_||_.
- The whole lipid molecule undergoes one-dimensional diffusion in the direction perpendicular to the membrane plane (in-out of the plane diffusion). This process is characterized by the force constant k, which characterizes the rigidity of the membrane in the normal direction of the membrane plane.
ii. The intermediate motions include the following three motions:

- The whole lipid molecule undergoes rotational diffusion around its long axis. This process is characterized by the distance between the long axis and the center of inertia of all the hydrogen atoms R_H_.
- The head group performs a head-flip-flop motion between the angles ±α with respect to the normal axis. The α values are 32.3° and 30° for DMPC and DMPG, respectively [23,24].
- The head group also performs a rotational motion around its own axis with the radius b_H_.
iii. The fast motions include the following two motions:

- The tail group undergoes two-dimensional diffusion within the membrane plane with its radius increasing as a function of the distance of the H atoms from the head group along the tail axis. This radius is defined as 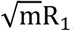, where R_1_ is the radius of the 2D-diffusion of H atoms closest to the head group and m denotes the index number of a methylene/methyl group of the tail group. The range of m is 1 ≤ m ≤ M, where M is the total number of methylene/methyl groups on the tail (M = 14 for both DMPC and DMPG).
- Hydrogen atoms in the methylene and methyl groups in both the head and tail groups undergo rotational jump-diffusion. Two-sites jump diffusion was assumed, where the residence times of the two sites are τ_1_ and τ_2_, respectively, with the jump distance corresponding to the H-C-H distance of d = 2.2 [Å] [25].

The above parameters R_||_, k, R_H_, b_H_, R_1_, and φ (= τ_1_/(τ_1_+τ_2_)), which characterize the geometry of motions of atoms in the lipids, were determined by the EISF/QISF analysis in the accompanying papers [21,22].

## 3. Results and discussion

### 3.1. Slow motions

The Q^2^-dependences of Γ_1_(Q), which corresponds to the slow motions, of all the samples are shown in Fig. 3. It is clearly seen that the Γ_1_(Q) values of all the samples do not reach zero as Q approaches 0.0 [Å^−1^] while they show asymptotic behavior at higher Q^2^ values, suggesting that the underlying process of this behavior is the jump diffusion within a confined space [14]. Note that in the previous study [19], the narrowest Lorentzian function of the DMPC MLB135 was interpreted as a continuous diffusion within a confined space. This discrepancy is due to the fact that the energy resolutions in the current study are much higher and the accessible Q-ranges are much larger than in the previous study, suggesting that the present data represent faster and smaller-scale motions. Therefore, the jump-diffusional behavior of Γ_1_(Q) indicates that each step of the global jump diffusion of the entire lipid molecule is resolved here. In the case of DMPC MLB45 at 283 K and 311 K, the Γ_1_(Q) values might be constant over the Q-range measured, suggesting that these motions are described by a diffusion on a linear segment or by rotational diffusion [26]. Since it is not possible to distinguish the type of motions due to the limited data points and large error bars, analysis was conducted for both types of motions for these two conditions.

**Figure 3.**
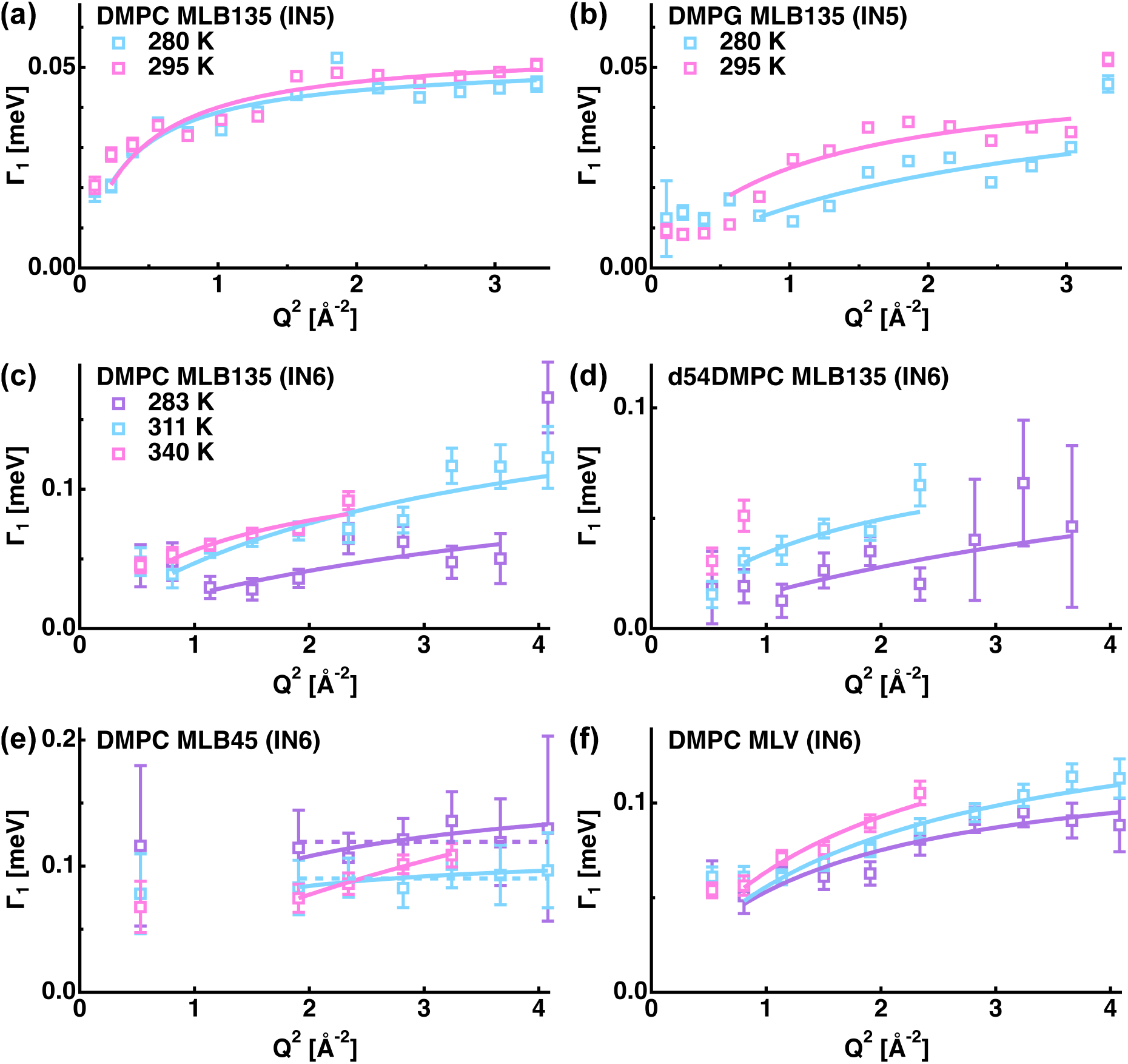
Q^2^-dependences of Γ_1_(Q) of DMPC MLB135 (a), DMPG MLB135 (b) measured on IN5, DMPC MLB135 (c), d54DMPC MLB135 (d), DMPC MLB45 (e), and DMPC MLV (f) measured on IN6. (a, b): The data points at 280 K and 295 K are shown in cyan and magenta, respectively. (c-f): The data points at 283 K, 311 K, and 340 K are shown in purple, cyan, and magenta, respectively. The solid lines show the corresponding fits by the equation describing the jump-diffusion (Eq. (2)) and the dotted lines (DMPC MLB45 at 283 K and 311 K) are the fits by the equation describing the diffusion on a linear segment (Γ_1_(Q) = 1/τ_1_, where τ_1_ is the correlation time.). Error bars are within symbols if not shown.

The radius of the confined space (*a*), which is related to the center-of-mass diffusion, was estimated from the Q_0_ value, at which Γ_1_(Q) begins to increase, based on the relation *a* = π/Q_0_ [27]. Since some data points were missing for DMPC MLB45 due to the experimental configuration, the *a* values were estimated for the other three samples, which were found to be 2.9–3.5 Å and 3.6–6.6 Å for IN6 and IN5, respectively, as shown in Table 2. The estimated *a* values of DMPC MLB135 are largely different between IN6 and IN5. One possibility of this difference would be the fact that it is not easy to determine the Q_0_ value due to the limited data points and fluctuations of the Γ_1_(Q) values in the low-Q region. Overall, the estimated *a* values here are smaller than the lateral distance between phospholipid molecules. However, since these values correspond to those resolved by the current time-windows, it is not surprising to obtain such smaller values.

**Table 2.** Estimated radii of the confined space *a* [Å] for slow motions.

|  | 280 K | 283 K | 295 K | 311 K | 340 K |
| --- | --- | --- | --- | --- | --- |
| DMPC MLB135 (IN6) |  | 2.9 |  | 2.9 | 2.9 |
| DMPC MLB135 (IN5) | 6.6 |  | 6.6 |  |  |
| d45DMPC MLB135 (IN6) |  | 3.5 |  | 3.5 | n/a |
| DMPC MLB45 (IN6) |  | n/a |  | n/a | n/a |
| DMPC MLV (IN6) |  | 3.5 |  | 3.5 | 3.5 |
| DMPG MLB135 (IN5) | 3.6 |  | 4.2 |  |  |

Next, dynamics’ parameters of the slow motions were estimated by the fitting of the Γ_1_(Q) values using the equation describing the jump diffusion [14];

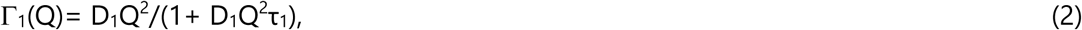

where D_1_ is the jump diffusion coefficient and τ_1_ is the residence time before the jump. Furthermore, based on these dynamics’ parameters, the jump distance l_1_ was also estimated by the relation I_1_ = √(4D_1_τ_1_) for in-plane motions, I_1_ = √(2D_1_τ_1_) for out-of-plane motions, and I_1_ = √(6D_1_τ_1_) for MLVs. In addition to this analysis, for DMPC MLB45 at 283 K and 311K, the correlation time was also calculated using the relation τ_1_ = 1/Γ_1_(Q) assuming the diffusion on a linear segment. However, the obtained τ_1_ values were the same within the errors as those obtained by the jump-diffusion model (Table S1). Therefore, in the following, discussion is made based on the dynamics’ parameters obtained by the jump-diffusion model. The summary of the obtained dynamics’ parameters is shown in Fig. 4 (the D_1_ and τ_1_ values are tabulated in Tables S1 and S2). The obtained parameters of DMPC MLB135 and DMPC MLB45 are found to be similar within the errors except at 340 K. This is in agreement with a MD simulation study showing that the center-of-mass diffusion coefficients of DMPC at 303 K are essentially the same between the in-plane and the out-of-plane directions (the difference is less than 10%) [28].

**Figure 4.**
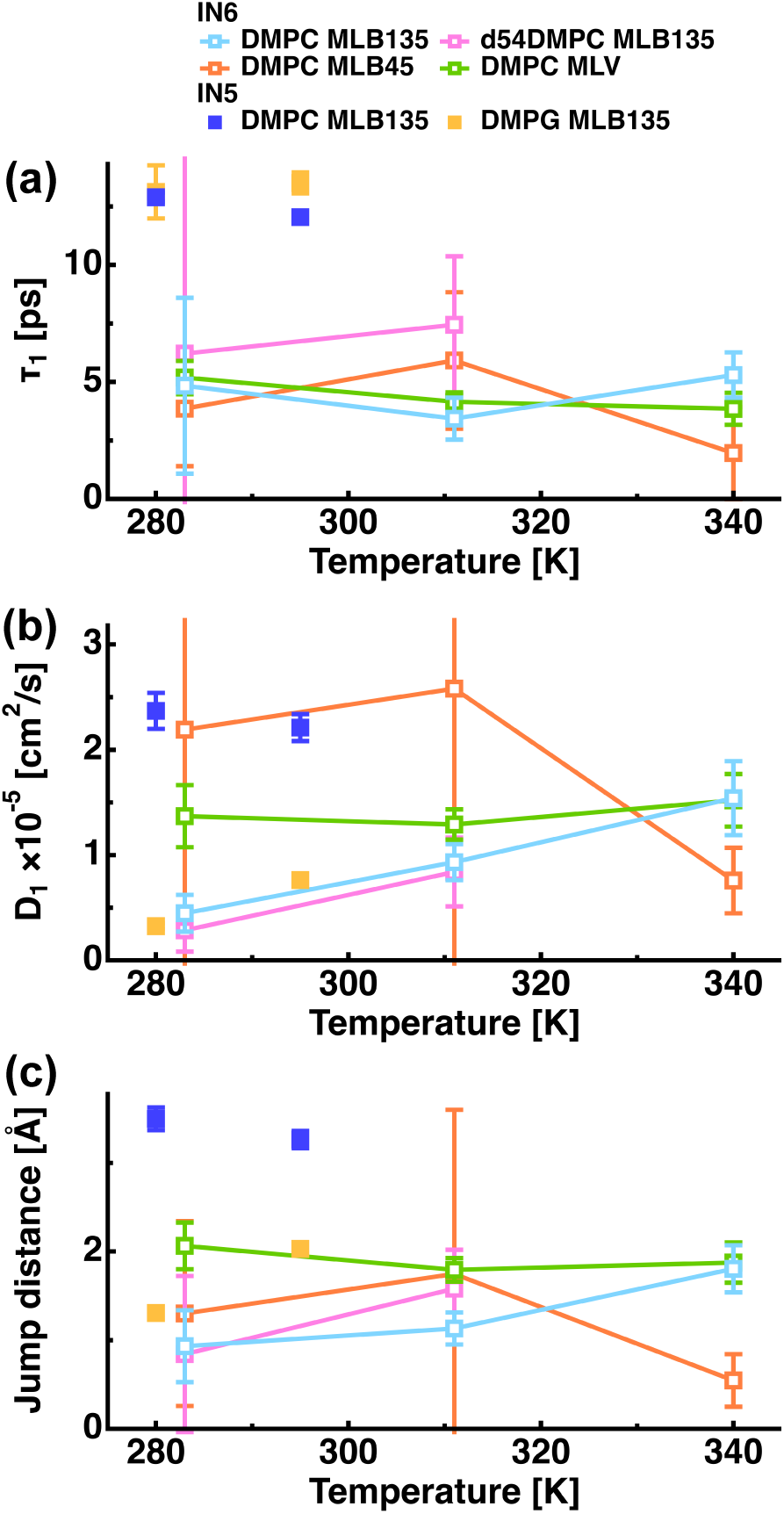
Temperature dependence of the residence time (τ_1_), the diffusion coefficient (D_1_), and the jump distance (I_1_) of the slow motions. The parameter values of DMPC MLB135, d54DMPC MLB135, DMPC MLB45, and DMPC MLV measured on IN6 are shown in cyan, magenta, orange, and green, respectively. Those of DMPC MLB135 and DMPG MLB135 measured on IN5 are shown in blue and pale orange, respectively.

When comparing the D_1_ values of d54DMPC MLB135 and DMPC MLB135, there is a tendency that D_1_ of the former is slightly smaller at both temperatures although these differences are within errors. The former mainly derives from the contribution of the head group and the latter contains contributions of both the head and the tail groups. This result thus implies that the diffusion of the head group might be slower than that of the tail group, which supports an assumption made in the Matryoshka-model, where the tail group motions are faster than the head group motions [21,22]. The tail group would undergo diffusive motions around the hinge between the tail and the head groups while the head group is less mobile, which increases the D_1_ value of the whole molecule compared with that of the head group as observed here. In a previous QENS study on the bilayers of dipalmitoyl-phosphatidylcholine (DPPC) with and without deuteration of its tail group, the same tendency has been observed [29]. The EISF/QISF analysis by the Matryoshka model on the same samples has shown that the diffusion radius of the head group (d54DMPC MLB135) on the membrane plane is larger than that of the whole molecule [21,22]. Together with this finding, the present results imply that the head group of DMPC takes more time to explore the available space than the tail group.

The dynamics’ parameters of the slow motions of DMPC MLB135 were found to be larger for IN5 than for IN6. The geometry of motions for the slow motion of DMPC MLB135 is also slightly different between the measurements on IN6 and those on IN5 [22]. Because MLBs measured on IN6 were more hydrated than those measured on IN5 (27 wt% and 10 wt% hydration for IN6 and IN5 measurements, respectively) and thus enhanced dynamics is expected for IN6, the differences in dynamics’ parameters between IN5 and IN6 observed here are not due to the difference in sample environments, but suggest that there is a distribution in the diffusive motions classified as the slow motions. The present results suggest that slight differences in atomic motions are detected by QENS and seen by application of the Matryoshka model, which proves their high sensitivity to decipher the dynamical nature of the samples.

The D_1_ values and the jump distance of DMPC MLB135 obtained on IN5 were found to be larger than those of DMPG MLB135, indicating that the diffusive motions are restrained in DMPG. Since the head group of DMPC has a larger mass than that of DMPG (the molecular weights are 104 and 92 Da for the head groups of DMPC and DMPG, respectively), it is expected that DMPC undergoes slower diffusive motions. However, the head group of DMPC possesses a positive charge, but is more hydrophobic than that of DMPG, which has two hydroxyl groups and thus polarity. It is likely that the hydration water molecules form less H-bonding with the head groups of DMPC than those of DMPG, indicating that the atoms in the head group of DMPC can move more freely. Because of these differences, the motions of the whole lipid molecule would also be enhanced for DMPC compared with DMPG.

Regarding the comparison between DMPC MLB135 and DMPC MLV on IN6, it is found that while the τ_1_ values are similar within the errors, the D_1_ value and the jump distance are larger for MLVs at 283 K and 311 K. This could be due to the fact that the MLVs are generally more hydrated than MLBs [13]. The present results are also consistent with the results of the EISF/QISF analysis using the Matryoshka model, where the radius of a circle within which the whole lipid molecule can diffuse is larger for MLVs than MLBs [22].

### 3.2. Intermediate motions

The Q^2^-dependences of Γ_2_(Q) of all the samples are shown in Fig. 5. The profiles of all the samples, except for DMPC MLB45, show non-zero values at low-Q regions, followed by an asymptotic behavior, suggesting that these motions are described by jump diffusion within a confined space [14]. On the other hand, the profiles for DMPC MLB45 do not show a Q^2^-dependent behavior, indicating that this motion is described by a diffusion on a linear segment or rotational diffusion [26]. Since the measurement of DMPC MLB45 mainly observes the motion perpendicular to the membrane plane, it is reasonable that only a one-dimensional motion is observed.

**Figure 5.**
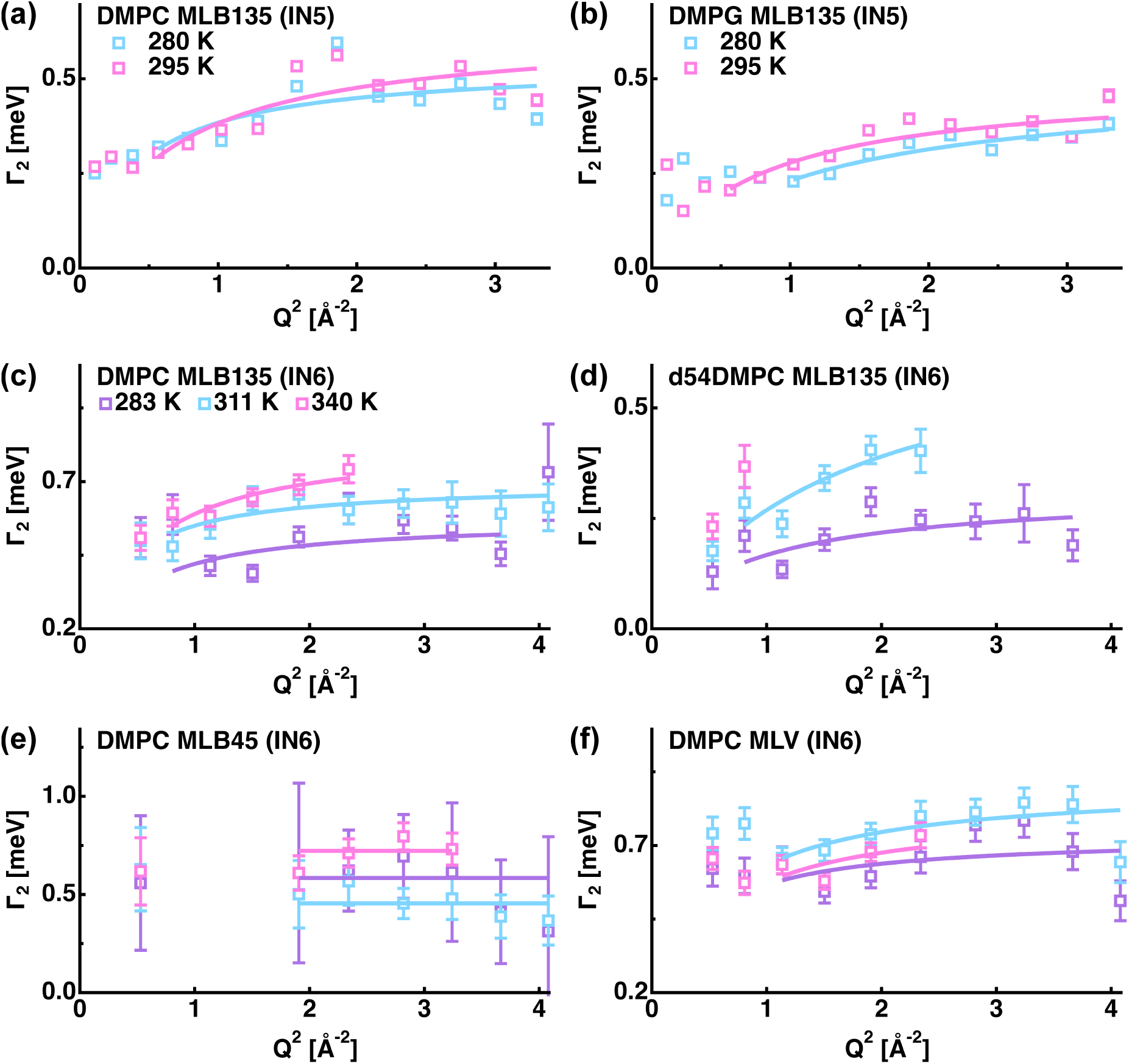
Q^2^-dependences of Γ_2_(Q) of the intermediate motions. The color scheme is the same as in Fig. 3. The solid lines show the fits by the equation describing the jump diffusion (Γ_2_(Q)= D_2_Q^2^/(1+ D_2_Q^2^τ_2_)) or by the equation describing the diffusion on a linear segment (Γ_2_(Q) = 1/τ_2_) for DMPC MLB45 (for the definition of the parameters, see the main text). Note that the scales on the ordinates are not the same in all the panels such that the fitting curves are better seen.

In order to obtain the dynamical parameters related to intermediate motions, the profiles in Fig. 5 were fitted by the equation describing the jump diffusion model, Γ_2_(Q)= D_2_Q^2^/(1+ D_2_Q^2^τ_2_), where D_2_ is the jump diffusion coefficient and τ_2_ is the residence time before the jump [14] in the Q^2^-range between 0.81 Å^−2^ and 4.08 Å^−2^ for the IN6 data and the Q^2^-range between 0.58 Å^−2^ and 3.30 Å^−2^ for the IN5 data. Furthermore, based on these dynamics’ parameters, the jump distance I_2_ was also estimated by the relation I_2_ = √(4D_2_τ_2_) for in-plane motions and I_2_ = √(6D_2_τ_2_) for MLVs. The correlation time of DMPC MLB45 was calculated using the relation τ_2_ = 1/Γ_2_(Q).

The residence time (correlation time), jump diffusion coefficient, and the jump distance parameters obtained by the above fits are shown in Fig. 6 (a), (b), and (c), respectively (the parameter values are tabulated in Tables S1 and S2). It is found that as the temperature increases, the τ_2_ values slightly decrease. There are no significant differences in the τ_2_ values between DMPC MLB135 and DMPC MLB45, suggesting that the degree of interaction between atoms is similar in directions both parallel and perpendicular to the membrane plane.

**Figure 6.**
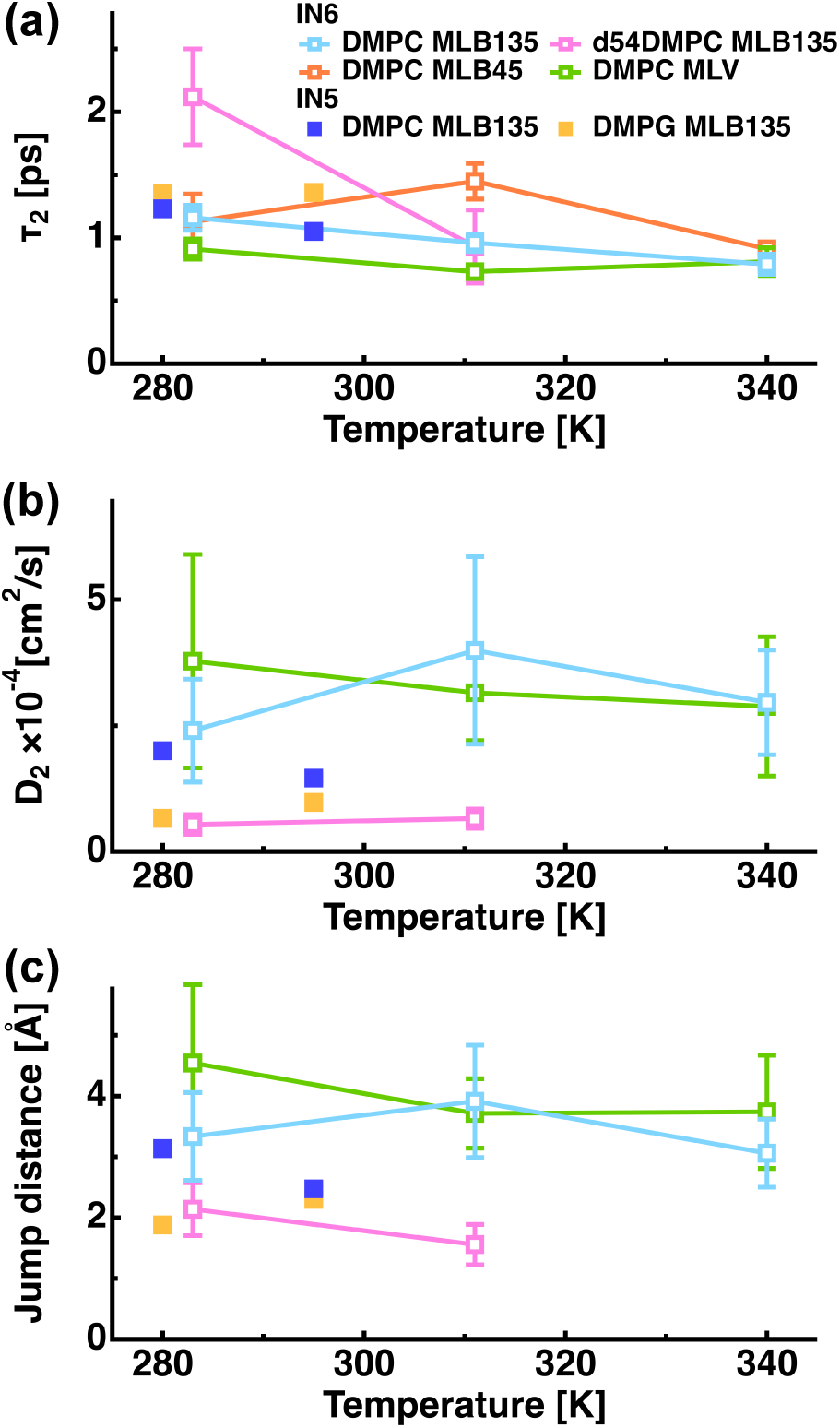
Temperature dependence of the residence time (τ_2_), the diffusion coefficient (D_2_), and the jump distance (I_2_) of the intermediate motions. The color scheme is the same as in Fig. 4.

The τ_2_ values of DMPC MLV were found to be smaller than those of DMPC MLB135, especially at 283 K and 311 K, while the D_2_ values and the l_2_ values were similar between them. This suggests that the atoms in MLVs move more rapidly than those in MLBs. Since lipid molecules are generally more hydrated in MLVs than in MLBs [13], it is likely that more hydration water enhances the frequency of the local motions in MLVs. Moreover, the mobile fraction of atoms, which is estimated by the Matryoshka model [22], is smaller for DMPC MLB135 than for DMPC MLV, suggesting that there exist more hydrogen atoms in MLBs that undergo slower motions than those in MLVs.

The D_2_ values shown in Fig. 6 (b) were larger than the translational diffusion coefficients of bulk water, which are 2.0−2.3 ×10^−5^ cm^2^/s at 293–300 K [30–32] although the residence times were larger than that of bulk water. The same phenomenon was previously reported for protein dynamics: A QENS study on human acetylcholinesterase (hAChE) has shown that the jump-diffusion coefficients of atoms in hAChE were lower or even higher than those of bulk water at 90 μeV and 50 μeV energy resolutions, respectively [33]. Since the time-scale and the length-scale of the observable motions depend on the energy resolution employed, the D_2_ values obtained in this study for atomic motions in lipids should be regarded as “effective” jump-diffusion coefficients, which depend on the instrumental energy resolution. It would thus be crucial to study the diffusive nature of local atomic motions in a wide range of time- and length-scale to obtain the holistic physical picture of the dynamical behavior of lipid molecules.

The D_2_ values of d54DMPC MLB135 were found to be much smaller than those of DMPC MLB135 while the τ_2_ values tend to be larger for the former, resulting in smaller l_2_ values for d54DMPC MLB135 than for DMPC MLB135. This suggests that the local atomic motions of the tails are faster than those of the head groups. This is in good agreement with the Matryoshka model [21], where it was necessary to ascribe the contribution of the tail groups to the fast motions in order to fully account for the EISFs and QISFs of the current samples. Since the Matryoshka model was established independently of the width analysis of Lorentzian functions described in this paper, this agreement further enhances the validity of the Matryoshka model. Moreover, it has been shown that the energetic barrier between conformational substates is higher for the head groups than for the tail groups [34], which is consistent with the present results.

The jump distances of the in-plane motions of the head group and whole molecule of DMPC were found to be ~2 Å and 3–4 Å, respectively. The latter value corresponds to the distance between the neighboring alkyl chains in the lipid molecules. On the other hand, according to the analysis based on the Matryoshka model, the radii within which the whole molecules can diffuse around are 1.8–2.4 Å (i.e. 3.6–4.8 Å in diameter) and 1.1–2.1 Å (2.2–4.2 Å in diameter) for the corresponding samples, respectively [22]. The comparison of these values suggests that while the whole DMPC molecule undergoes a jump motion over a distance corresponding to the neighboring alkyl chains, its head group moves by about half of the maximum distance available for it. This behavior of the head group could be due to the head-flip-flop motion as introduced in the intermediate motion of the Matryoshka model, which changes the distance between the centers of mass of the head group and the tail groups projected onto the membrane plane. Furthermore, the current study suggests that all of the above motions take place during the timescale of 1–2 ps on average.

As for DMPC MLB135, all of the three parameter values obtained from the IN5 data were similar to those obtained from the IN6 data, showing that the identical motions are resolved at both energy resolutions unlike the slow motions described above. When comparing the parameters between DMPC MLB135 and DMPG MLB135 on IN5, the residence time of DMPC was smaller than that of DMPG while the jump diffusion coefficient of DMPC is larger, resulting in a slightly larger jump distance for DMPC. This indicates that the local motions are enhanced for DMPC compared to DMPG, which is in agreement with the findings for the slow motions. The reduction in local dynamics of DMPG MLB135 compared with DMPC MLB135 is consistent with the results obtained from the EISF/QISF analysis by the Matryoshka model where the radius of the two-dimensional diffusion of the tail group on the membrane plane is smaller for DMPG [22].

### 3.3. Fast motions

As shown in Fig. 7 (a-f), the widths of the broadest Lorentzian functions corresponding to the fast motions were found to be independent of Q for all the samples, suggesting that these motions are mainly described by rotational jump diffusion [14] as seen in hydrogen atoms in the methylene and methyl groups. On the other hand, the Matryoshka model assumes another kind of motions in addition to the rotational jump diffusion for the fast motions, which is the two-dimensional diffusion of the hydrogen atoms in the tail groups within the membrane plane. The width of the Lorentzian function describing this motion would increase as the Q^2^ values rise, which is not observed for Γ_3_(Q) here, suggesting that the contribution of the rotational jump diffusion to Γ_3_(Q) is dominant. This is in line with the result of the analysis based on the Matryoshka model in that the integrated value of the A_jump_(Q) over the Q-range employed is much larger than that of A_tail_(Q) [21].

**Figure 7.**
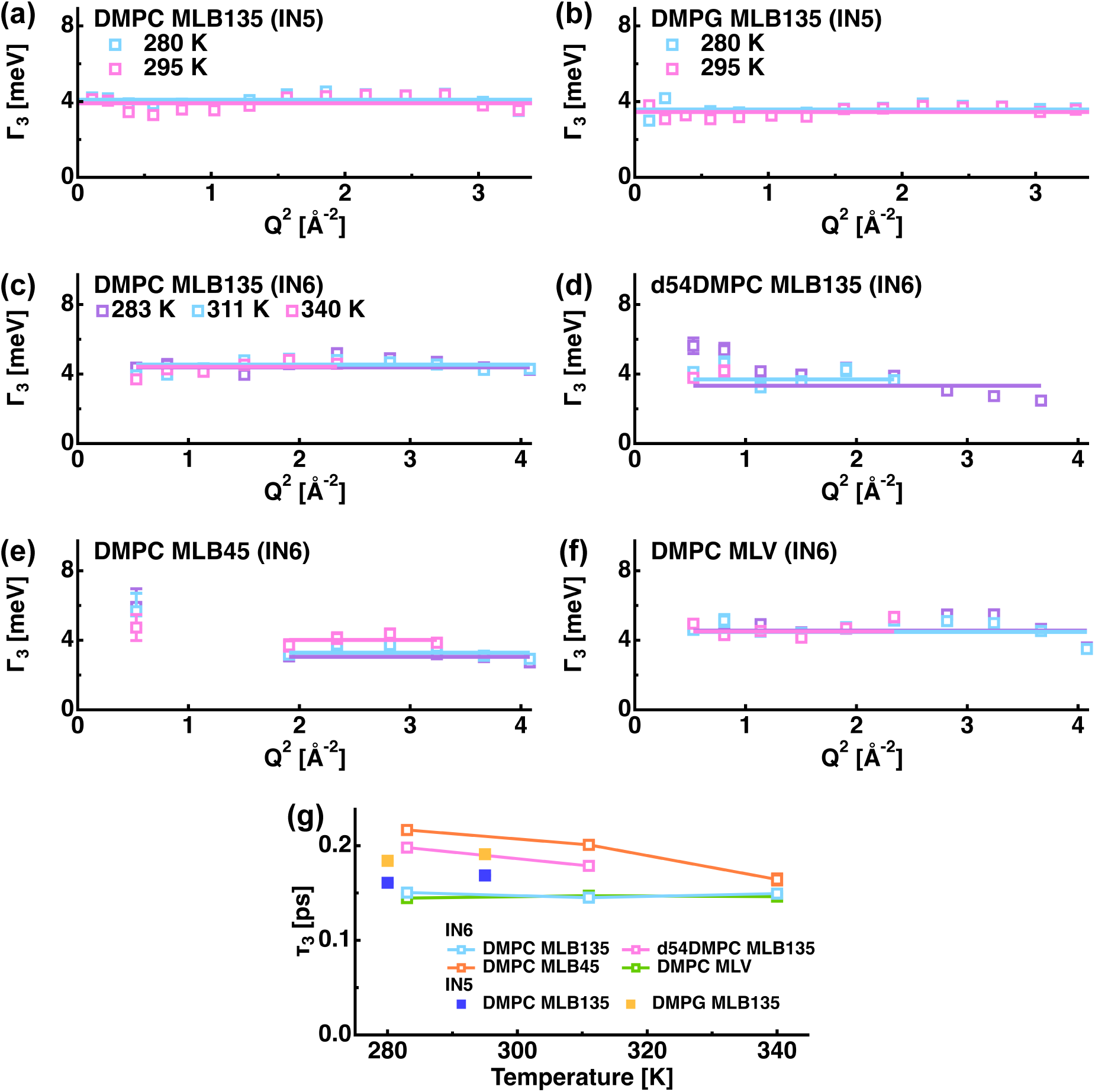
Summary of the parameters obtained for the fast motions. (a-f): Q^2^-dependences of Γ_3_(Q). The color scheme is the same as in Figs. 3 and 5. The solid lines show the corresponding fits by the equation describing the rotational jump diffusion (Γ_3_(Q) = 1/τ_3_) (see also the text). (g): Temperature dependence of the correlation time (τ_3_) of the fast motions. The color scheme is the same as in Figs. 4 and 6.

The correlation time of these motions was calculated from the relation τ_3_ = 1/Γ_3_(Q), and its dependence on the temperature is shown in Fig. 7 (g) (the τ_3_ values are tabulated in Tables S1 and S2). The τ_3_ values of DMPC MLB135 on IN6 and DMPC MLV are found to be independent of the temperature and to be the same within the errors, suggesting that the frequency of the methyl and methylene rotations is independent of the local environments in which they reside (i.e. in the MLB or in the MLV). The τ_3_ values of d54DMPC MLB135 are larger than those of DMPC MLB135, indicating that the rotational jump diffusion is enhanced for the tail group compared with for the head group, which is consistent with the basic assumption of the Matryoshka model that the tail motions are faster than the head group motions [21,22]. The τ_3_ values of DMPC MLB45 are found to be larger than those of DMPC MLB135, implying that the fast motions show anisotropy (i.e. parallel or perpendicular to the membrane plane).

The τ_3_ values of DMPC MLB135 measured on IN5 were found to be slightly smaller than those of DMPG MLB135, suggesting that the frequency of the rotational jump diffusion of the methyl and methylene groups is higher for DMPC MLB135, which is consistent with the enhanced dynamics of DMPC MLB135 for the slow and the intermediate motions. Furthermore, the τ_3_ values of DMPC MLB135 were similar between IN5 and IN6, suggesting that the same type of motions is observed on both spectrometers as the intermediate motions. It thus appears that the different hydration state of the DMPC MLB135 between IN5 and IN6 as described above does not significantly affect the intermediate and fast motions. In the Matryoshka model, it was necessary to include the contribution of the rotational jump diffusion in the methyl and methylene groups in order to describe the QISF of the fast motion [21,22]. The present results indicate that the rotational jump in these chemical groups occurs every 0.2 ps.

### 3.4 Comparison of Γ_1_(Q), Γ_2_(Q), and Γ_3_(Q) between experiments and Matryoshka model

All of the above analyses were carried out independently of the application of the Matryoshka model and despite this fact, the present findings support the results obtained by the EISF/QISF analysis using this model [21,22]. In order to see further whether the Matryoshka model can reproduce the energy widths of the QENS spectra, the experimental Γ_1_(Q), Γ_2_(Q), and Γ_3_(Q) values were compared with those based on the Matryoshka model. Analytical equations of Γ_1_(Q), Γ_2_(Q), and Γ_3_(Q) based on this model, which are given in the Supplementary material, were fitted to the measured corresponding profiles. Note that only the samples other than MLVs were used for the fitting, because the derivation of the analytical expressions of Γ_1_(Q), Γ_2_(Q), and Γ_3_(Q) for MLVs is not straightforward. In the fitting, the values of the parameters (R_||_, k, R_H_, b_H_, R_1_, and φ), which were determined by the EISF/QISF analysis [22], were fixed except for the fitting of Γ_2_(Q) with the following cases:

- R_H_ of DMPC MLB135 at 283 K on IN6
- R_1_ of DMPC MLB135 at 340 K on IN6
- b_H_ of d54DMPC MLB135 at 283 K and 311 K on IN6
- b_H_ of DMPC MLB135 and DMPG MLB135 at 280 K and 295 K on IN5

The above parameters were allowed to change during the fitting because of their large errors estimated by the EISF/QISF analysis [22]. The fitting results are shown in Fig. 8 (the values of each parameter are tabulated in Tables S3 and S4). Although in some cases, the above parameter values were outside the range determined at the corresponding temperature by the EISF/QISF analysis [22] (these values are shown with asterisk in Tables S3 and S4), all of these parameter values were still within the range determined at other temperatures, suggesting that these values are not physically unrealistic ones. From Fig. 8, it is seen that the resultant fitting curves are able to reproduce the experimental Γ_1_(Q), Γ_2_(Q), and Γ_3_(Q) values quite well regardless of the instruments. Regarding the Γ_1_(Q) of in-plane motions, i.e. DMPC MLB135, DMPG MLB135, and d54DMPC MLB135, as already discussed in the section 3.1 (see also Fig. 3), because of the wide Q-range and the low energy resolutions, our experimental data appear to permit to see the discrete steps of the global diffusion of the whole lipid molecules. Thus, description of such detailed motions, which are independent of EISF/QISFs, would improve further the quality of the fitting. All in all, the Matryoshka model can reproduce not only the EISF/QISF profiles, but also the energy widths of the QENS spectra, enhancing the reliability of this model to describe the lipid dynamics.

**Figure 8.**
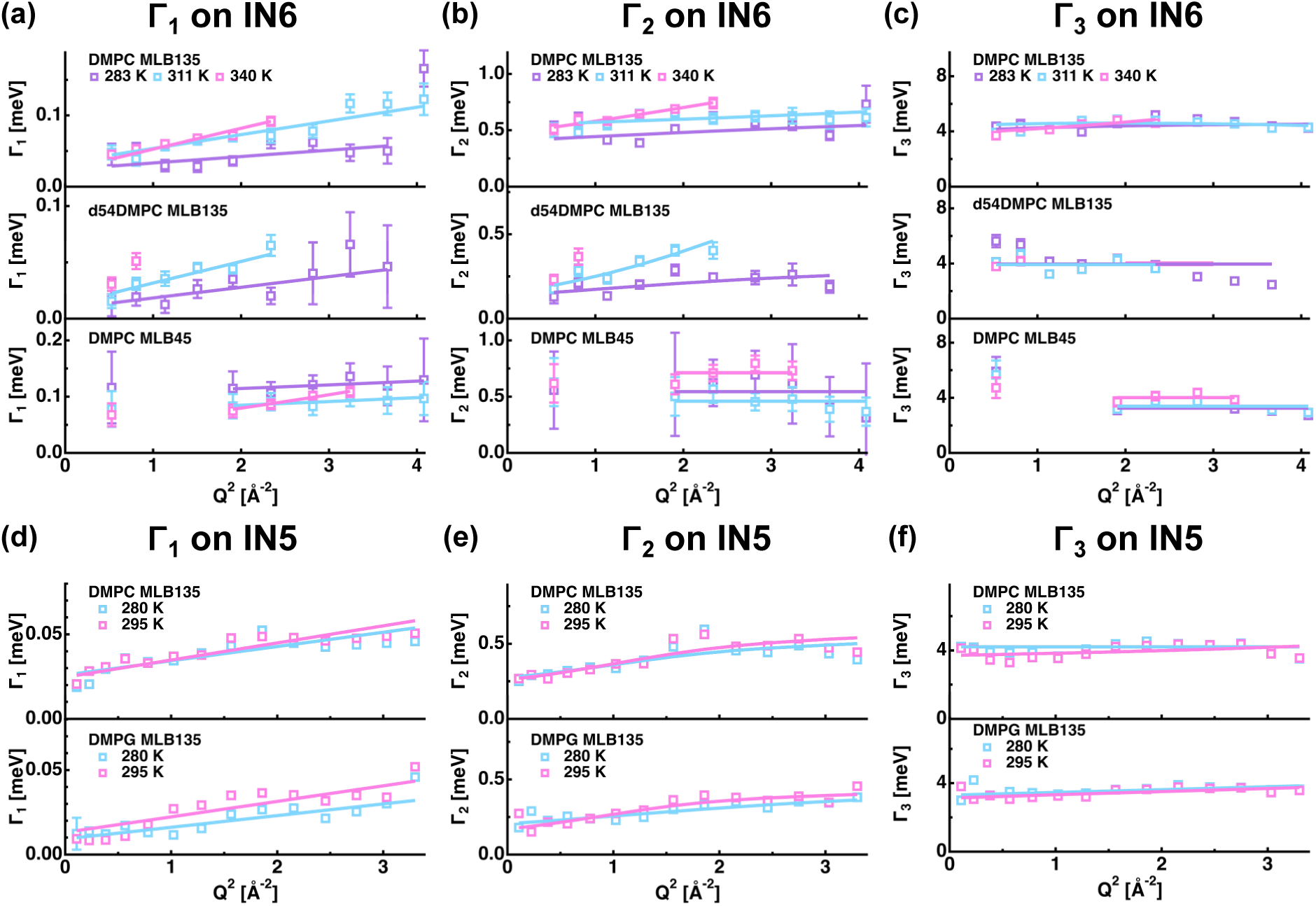
Comparison of the experimental Γ_1_(Q), Γ_2_(Q), and Γ_3_(Q) profiles with those predicted by the Matryoshka model. Q^2^-dependences of Γ_1_(Q), Γ_2_(Q), and Γ_3_(Q) on IN6 are shown in (a), (b), and (c), respectively. The data points at 283 K, 311 K, and 340 K are shown in purple, cyan, and magenta, respectively. (d), (e), and (f) show the Q^2^-dependences of Γ_1_(Q), Γ_2_(Q), and Γ_3_(Q) on IN5, respectively. The data points at 280 K and 295 K are shown in cyan and magenta, respectively. The solid lines show the corresponding fits based on the Matryoshka model (Eq. (1-12) in the Supplementary material).

## 4. Concluding remarks

In this study, the diffusive nature of the motions contained in the QENS spectra of DMPC and DMPG in various forms was analyzed and thus relaxational aspects of the Matryoshka model were characterized. The three Lorentzian functions, which were obtained by decomposing the QENS spectra, showed significantly distinctive relaxational features with their widths spanning about two orders of magnitude. Regardless of this variety, the Matryoshka model can satisfactorily describe the various motions contained in each of these components. The current results of the analysis of the widths of these Lorentzian functions, which was carried out independently of the development of the Matryoshka model, not only provides information on the relaxational aspects of the motions contained in this model, but also corroborates the parameter values such as the amplitudes and the mobile fractions of atomic motions obtained by the EISF/QISF analysis using the Matryoshka model, which enhances the validity of the model. The Matryoshka model can thus serve as a powerful tool to decipher the intramolecular motions of lipid molecules and demonstrate its power in characterizations of more complex systems such as the mixtures of different kinds of lipids, protein-lipid complexes, and eventually natural cell membranes in the near future.

## Acknowledgements

The authors thank the Institut Laue-Langevin for the allocation of the beam time to perform the experiments. AC is supported by the JP Aguilar scholarship from the Foundation CFM for her PhD thesis.

## Supplementary material

Parameter values obtained from the QENS spectra, analytical expressions of Γ_1_(Q), Γ_2_(Q), and Γ_3_(Q) based on the Matryoshka model are found in the Supplementary material.

