## Supplementary material for "The dynamical Matryoshka model: 3. Diffusive nature of the atomic motions contained in a new dynamical model for deciphering local lipid dynamics"

[f]Instit. Univ. de France

### *1. Analytical expressions of $\Gamma_1(Q)$, $\Gamma_2(Q)$, and $\Gamma_3(Q)$*

The half-width at half-maximum (HWHM) of the slow motion $\Gamma_1(Q)$ is represented as [1]

$$\Gamma_1 = \frac{(1-A_{in-out})\Gamma_{in-out} + (1-A_{2d})\Gamma_{2d}}{1- A_{in-out}A_{2d}}, \ (1)$$

where $A_{in-out}$ and $\Gamma_{in-out}$ denote the elastic incoherent structure factor (EISF) and the HWHM of the in-out-of plane diffusion of the whole lipid molecule, respectively. $A_{2d}$ and $\Gamma_{2d}$ denote the EISF and the HWHM of the 2D-diffusion of the whole lipid molecule, respectively. $A_{in-out}$, $A_{2d}$, $\Gamma_{in-out}$, and $\Gamma_{2d}$ are expressed as

$$A_{in-out} = \exp\left\{-(Q\cos\gamma)^2\left(\frac{k_B T}{k}\right)\right\}, \ (2)$$

$$A_{2d} = \left\{\frac{2J_1(QR_{||}\sin\gamma)}{QR_{||}\sin\gamma}\right\}^2, \ (3)$$

$$\Gamma_{in-out} = \left\{(Q\cos\gamma)^2\left(\frac{k_B T}{k}\right)\frac{1}{\tau_{in-out}}\right\} / \left[1-\exp\left\{-(Q\cos\gamma)^2\left(\frac{k_B T}{k}\right)\right\}\right], \ (4)$$

$$\Gamma_{2d} = \left\{\frac{(x_0^1)^2}{R_{||}^2} + Q^2\right\}D_{||}, \ (5)$$

where $\gamma$ is the angle between the scattering vector ($\vec{Q}$) and the normal to the membrane plane, i.e. $\gamma = 0$ and $\gamma = \frac{\pi}{2}$ refer to the direction of $\vec{Q}$ perpendicular and parallel to the membrane plane, respectively. $k_B$ is the Boltzmann constant, T is the temperature. $k$ and $\tau_{in-out}$ are the force constant and the correlation time of the in-out of the plane motions of the whole lipid molecule. $R_{||}$ and $D_{||}$ denote the radius and the translational diffusion coefficient of the 2D-diffusion of the whole lipid molecule, respectively. $x_0^1$ is equal to 1.84118.

The HWHM of the intermediate motion $\Gamma_2(Q)$ is represented as

$$\Gamma_2 = \frac{z(1-A_{head})\Gamma_{head} + [z+(1-z)A_{tail}](1-A_{rot})\Gamma_{rot}}{z(1- A_{head}A_{rot}) + (1-z)A_{tail}(1-A_{rot})}, \ (6)$$

where $A_{head}$ and $\Gamma_{head}$ denote the EISF and the HWHM of the motions of the head group, respectively. $A_{tail}$ and $\Gamma_{tail}$ denote the EISF and the HWHM of the motions of the tail group, respectively. $A_{rot}$ denotes the EISF of the rotational diffusion of the whole lipid molecule around its long axis.

$A_{head}$, $A_{tail}$, $\Gamma_{head}$, $\Gamma_{tail}$, and $\Gamma_{rot}$ are expressed as

$$A_{head} = \left[\sum_{l=-\infty}^{+\infty} J_{2l}(Qb_H\cos\gamma\sin\alpha)J_l\left(Qb_H\sin\gamma\cos^2\frac{\alpha}{2}\right)\right]^2, \ (7)$$

$$A_{tail} = \frac{1}{M}\sum_{m=1}^{M}\left[\frac{2J_1(QR_m\sin\gamma)}{QR_m\sin\gamma}\right]^2, \ (8)$$

$$\Gamma_{head} = D_{head} + \frac{1}{\tau_{ff}}, \ (9)$$

$$\Gamma_{tail} = D_{tail}\left[\frac{1}{M}\sum_{m=1}^{M}(1-A_{tail})\left\{\frac{(x_0^1)^2}{R_m^2} + Q^2\right\}\right] / \left[\frac{1}{M}\sum_{m=1}^{M}(1-A_{tail})\right], \ (10)$$

$\Gamma_{\mathrm{rot}} = \mathrm{D}_{\mathrm{rot}}$, (11)

where $D_{head}$ and $\tau_{ff}$ are the rotational diffusion coefficient and the correlation time of the flip-flop motion of the head group, respectively. $D_{tail}$ is the translational diffusion coefficient of the 2D-diffusion of the tail group. $D_{rot}$ is the rotational diffusion coefficient of the whole lipid molecule.

Finally, the HWHM of the fast motion $\Gamma_3(Q)$ is represented as

$$\Gamma_3 = \frac{(1-\mathrm{A_{jump}})\Gamma_{\mathrm{jump}} + (1-\mathrm{z})(1-\mathrm{A_{tail}})\Gamma_{\mathrm{tail}}}{\mathrm{z}(1-\mathrm{A_{jump}}) + (1-z)(1-\mathrm{A_{jump}}\mathrm{A_{tail}})}, \quad (12)$$

where $\Gamma_{\mathrm{jump}}$ denotes the HWHM of the rotational jump diffusion of the methylene/methyl groups.

### *2. Summary of the dynamics parameters obtained in this study*

Tables S1 and S2 below show the list of parameter values obtained in this study. Tables S3 and S4 show the list of parameter values obtained by the fitting of the experimental data by the Matryoshka-model. In all tables, the values in the parentheses are error values associated with the fitting. Note that, in Tables S3 and S4, the parameter values, which are outside the range determined by the EISF/QISF analysis [2], are shown with asterisk (*).

**Table S1. Summary of the parameter values obtained from the energy widths of the QENS spectra measured on IN6.**

| DMPC MLB135 | | | |
|---|---|---|---|
| Parameters | 283 K | 311 K | 340 K |
| $\tau_1$ [ps] | 5 (4) | 3.4 (0.9) | 5 (1) |
| $D_1$ [$\times 10^{-6}$ cm$^2$/s] | 4 (2) | 9 (2) | 15.4 (0.4) |
| $l_1$ [Å] | 0.9 (0.4) | 1.1 (0.2) | 1.8 (0.3) |
| $\tau_2$ [ps] | 1.2 (0.1) | 0.96 (0.07) | 0.79 (0.08) |
| $D_2$ [$\times 10^{-4}$ cm$^2$/s] | 2 (1) | 4 (2) | 3 (1) |
| $l_2$ [Å] | 3.3 (0.7) | 3.9 (0.9) | 3.0 (0.6) |
| $\tau_3$ [ps] | 0.150 (0.002) | 0.145 (0.002) | 0.149 (0.002) |

| DMPC MLB45 | | | |
|---|---|---|---|
| Parameters | 283 K | 311 K | 340 K |
| $\tau_1$ [ps] (jump diffusion) | 4 (2) | 6 (3) | 2 (2) |
| $\tau_1$ [ps] (diffusion on a linear segement) | 5.5 (0.5) | 7.3 (0.6) | n.d. |
| $D_1$ [$\times 10^{-6}$ cm$^2$/s] | 22 (32) | 26 (53) | 8 (3) |
| $l_1$ [Å] | 2 (1) | 2 (3) | 0.8 (0.4) |
| $\tau_2$ [ps] | 1.1 (0.2) | 1.4 (0.1) | 0.91 (0.05) |
| $\tau_3$ [ps] | 0.216 (0.005) | 0.201 (0.004) | 0.164 (0.006) |

| d54DMPC MLB135 | | | |
|---|---|---|---|
| Parameters | 283 K | 311 K | 340 K |
| $\tau_1$ [ps] | 6 (12) | 7 (3) | n.d. |
| $D_1$ [$\times 10^{-6}$ cm$^2$/s] | 3 (2) | 8 (3) | n.d. |
| $l_1$ [Å] | 0.8 (0.9) | 1.6 (0.4) | n.d. |
| $\tau_2$ [ps] | 2.1 (0.4) | 0.9 (0.3) | n.d. |

| | | | |
|---|---|---|---|
| $D_2$ [×10$^{-4}$ cm$^2$/s] | 0.5 (0.2) | 0.6 (0.2) | n.d. |
| $l_2$ [Å] | 2.1 (0.4) | 1.6 (0.3) | n.d. |
| $\tau_3$ [ps] | 0.198 (0.004) | 0.179 (0.005) | n.d. |

The parameters at 340 K were not determined due to the limited data points.

| | DMPC MLV | | |
|---|---|---|---|
| Parameters | 283 K | 311 K | 340 K |
| $\tau_1$ [ps] | 5.2 (0.7) | 4.2 (0.4) | 3.8 (0.7) |
| $D_1$ [×10$^{-6}$ cm$^2$/s] | 13.7 (0.3) | 12.9 (0.1) | 15.2 (0.2) |
| $l_1$ [Å] | 1.7 (0.2) | 1.5 (0.1) | 1.5 (0.2) |
| $\tau_2$ [ps] | 0.91 (0.08) | 0.73 (0.05) | 0.8 (0.1) |
| $D_2$ [×10$^{-4}$ cm$^2$/s] | 4 (2) | 3.2 (0.9) | 3 (1) |
| $l_2$ [Å] | 4 (1) | 3.0 (0.5) | 3.0 (0.8) |
| $\tau_3$ [ps] | 0.145 (0.002) | 0.147 (0.002) | 0.146 (0.003) |

**Table S2. Summary of the parameter values obtained from the energy widths of the QENS spectra measured on IN5.**

| | DMPC MLB135 | | DMPG MLB135 | |
|---|---|---|---|---|
| Parameters | 280 K | 295 K | 280 K | 295 K |
| $\tau_1$ [ps] | 12.9 (0.2) | 12.0 (0.2) | 13 (1) | 13.5 (0.4) |
| $D_1$ [×10$^{-6}$ cm$^2$/s] | 24 (2) | 22 (1) | 3.2 (0.3) | 7.6 (0.5) |
| $l_1$ [Å] | 3.5 (0.1) | 3.3 (0.1) | 1.30 (0.08) | 2.03 (0.07) |
| $\tau_2$ [ps] | 1.23 (0.02) | 1.05 (0.02) | 1.35 (0.04) | 1.36 (0.02) |
| $D_2$ [×10$^{-4}$ cm$^2$/s] | 2.0 (0.1) | 1.46 (0.06) | 0.66 (0.03) | 0.97 (0.03) |
| $l_2$ [Å] | 3.1 (0.1) | 2.48 (0.06) | 1.88 (0.05) | 2.30 (0.04) |
| $\tau_3$ [ps] | 0.1608 (0.0004) | 0.1686 (0.0004) | 0.1840 (0.0004) | 0.1909 (0.0004) |

**Table S3. Summary of the parameter values obtained by fitting the $\Gamma_1(Q)$, $\Gamma_2(Q)$, and $\Gamma_3(Q)$ of the Matryoshka-model to the corresponding profiles measured on IN6.**

| DMPC MLB135 | | | |
|---|---|---|---|
| Parameters | 283 K | 311 K | 340 K |
| $D_{\|\|}$ [$\times 10^{-6}$ cm$^2$/s] | 1.4 (0.1) | 2.9 (0.1) | 4.4 (0.1) |
| $\Gamma_{head}$ [meV] | 1 (1) | 0.6 (0.3) | 2 (26) |
| $D_{rot}$ [$\times 10^{11}$ /s] | 1 (37) | 8 (3) | 1 (96) |
| $R_1$ [Å] | 0.43 | 0.42 | 0.44 (3.6)* |
| $R_H$ [Å] | 0.4 (0.4) | 0.46 | 0.62 |
| $D_{tail}$ [$\times 10^{-5}$ cm$^2$/s] | 14 (3) | 7 (2) | 6 (2) |
| $\tau_1$ [ps] | 0.2 (0.1) | 0.07 (0.01) | 0.36 (0.07) |
| $\tau_2$ [ps] | 2 (2) | 0.9 (0.2) | 1.0 (0.2) |

The $R_1$ values at 283 K and 311 K and the $R_{HL}$values at 311 K and 340 K were fixed to the corresponding values obtained by EISF/QISF analysis [2]. $D_{||}$ is obtained by Eq. (5).

| DMPC MLB45 | | | |
|---|---|---|---|
| Parameters | 283 K | 311 K | 340 K |
| $\tau_{in-out}$ [ps] | 6.4 (0.5) | 9.0 (0.8) | 15.9 (0.7) |
| $\Gamma_{head}$ [meV] | 0.5 (0.1) | 0.46 (0.07) | 0.71 (0.08) |
| $\tau_1$ [ps] | 0.27 (0.04) | 0.26 (0.04) | 0.22 (0.03) |
| $\tau_2$ [ps] | 0.8 (0.1) | 0.8 (0.1) | 0.63 (0.08) |

| d54DMPC MLB135 | | | |
|---|---|---|---|
| Parameters | 283 K | 311 K | 340 K |
| $D_{\|\|}$ [$\times 10^{-6}$ cm$^2$/s] | 1.4 (0.2) | 2.9 (0.2) | n.d. |
| $\Gamma_{head}$ [meV] | 0.13 (0.03) | 0.0 (0.9) | n.d. |
| $b_H$ [Å] | 1.85 (3.46) | 1.77 (3.19)* | n.d. |
| $D_{rot}$ [$\times 10^{11}$ /s] | 3 (1) | 35 (76) | n.d. |
| $\tau_1$ [ps] | 0.23 (0.03) | 0.20 (0.03) | n.d. |
| $\tau_2$ [ps] | 0.57 (0.07) | 1.1 (0.1) | n.d. |

The parameters at 340 K are not determined due to the limited data points. $D_{||}$ is obtained by Eq. (5).

**Table S4. Summary of the parameter values obtained by fitting the Γ$_1$(Q), Γ$_2$(Q), and Γ$_3$(Q) of the Matryoshka-model to the corresponding profiles measured on IN5.**

| | DMPC MLB135 | | DMPG MLB135 | |
|---|---|---|---|---|
| Parameters | 280 K | 295 K | 280 K | 295 K |
| $D_{||}$ [×10$^{-6}$ cm$^2$/s] | 1.281 (0.009) | 1.539 (0.008) | 1.05 (0.01) | 1.39 (0.01) |
| $\Gamma_{head}$ [meV] | 0.215 (0.008) | 0.212 (0.006) | 0.12 (0.04) | 0.112 (0.005) |
| $b_H$ [Å] | 2.8 (0.1)* | 2.8 (0.1) | 1.9 (0.3)* | 2.8 (0.1)* |
| $D_{rot}$ [×10$^{11}$ /s] | 11.99 (0.03) | 10.40 (0.01) | 6.9 (0.7) | 7.72 (0.09) |
| $R_H$ [Å] | 0.44* | 0.44 | 0.43 | 0.46 |
| $D_{tail}$ [×10$^{-5}$ cm$^2$/s] | n.d. | 2.2 (0.1) | 2.5 (0.1) | 3.0 (0.1) |
| $\tau_1$ [ps] | n.d. | 0.303 (0.003) | 0.312 (0.004) | 0.321 (0.004) |
| $\tau_2$ [ps] | n.d. | 0.588 (0.006) | 0.664 (0.008) | 0.75 (0.01) |

The $D_{tail}$, $\tau_1$ and $\tau_2$ values of DMPC MLB135 at 280 K cannot be determined because of $A_{jump} = 1$ in Eq. (12). The $R_H$ value of DMPC MLB135 at 280 K was fixed to 0.44 Å instead of 0.01 Å [2] to obtain a better fitting, and The $R_H$ values of other conditions were fixed to the corresponding values obtained by EISF/QISF analysis [2]. $D_{||}$ is obtained by Eq. (5).

[2] A. Cissé, T. Matsuo, M. Plazanet, F. Natali, M.M. Koza, J. Olivier, D.J. Bicout, J. Peters, The dynamical Matryoshka model: 2 A new modelling of local lipid dynamics at the sub-nanosecond timescale in phospholipid membranes, BBA - Biomembr., (n.d.).
